# OncoOmics approaches to reveal essential genes in breast cancer: a panoramic view from pathogenesis to precision medicine

**DOI:** 10.1101/638866

**Authors:** Andrés López-Cortés, César Paz-y-Miño, Santiago Guerrero, Alejandro Cabrera-Andrade, Stephen J. Barigye, Cristian R. Munteanu, Humberto González-Díaz, Alejandro Pazos, Yunierkis Pérez-Castillo, Eduardo Tejera

**Affiliations:** Centro de Investigación Genética y Genómica, Facultad de Ciencias de la Salud Eugenio Espejo, Universidad UTE, Mariscal Sucre Avenue, Quito 170129, Ecuador; RNASA-IMEDIR, Computer Science Faculty, University of Coruna, Coruna 15071, Spain; Carrera de Enfermería, Facultad de Ciencias de la Salud, Universidad de Las Américas, Avenue de los Granados, Quito 170125, Ecuador; Grupo de Bio-Quimioinformática, Universidad de Las Américas, Avenue de los Granados, Quito 170125, Ecuador; Department of Chemistry, McGill University, 801 Sherbrooke Street West, Montreal, QC H3A 0B8, Canada; INIBIC, Institute of Biomedical Research, CHUAC, UDC, Coruna 15006, Spain; Department of Organic Chemistry II, University of the Basque Country UPV/EHU, Leioa 48940, Biscay, Spain; IKERBASQUE, Basque Foundation for Science, Bilbao 48011, Biscay, Spain; Escuela de Ciencias Físicas y Matemáticas, Universidad de Las Américas, Avenue de los Granados, Quito 170125, Ecuador; Facultad de Ingeniería y Ciencias Agropecuarias, Universidad de Las Américas, Avenue de los Granados, Quito 170125, Ecuador

## Abstract

Breast cancer (BC) is a heterogeneous disease where each OncoOmics approach needs to be fully understood as a part of a complex network. Therefore, the main objective of this study was to analyze genetic alterations, signaling pathways, protein-protein interaction networks, protein expression, dependency maps and enrichment maps in 230 previously prioritized genes by the Consensus Strategy, the Pan-Cancer Atlas, the Pharmacogenomics Knowledgebase and the Cancer Genome Interpreter, in order to reveal essential genes to accelerate the development of precision medicine in BC. The OncoOmics essential genes were rationally filtered to 144, 48 (33%) of which were hallmarks of cancer and 20 (14%) were significant in at least three OncoOmics approaches: RAC1, AKT1 CCND1, PIK3CA, ERBB2, CDH1, MAPK14, TP53, MAPK1, SRC, RAC3, PLCG1, GRB2, MED1, TOP2A, GATA3, BCL2, CTNNB1, EGFR and CDK2. According to the Open Targets Platform, there are 111 drugs that are currently being analyzed in 3151 clinical trials in 39 genes. Lastly, there are more than 800 clinical annotations associated with 94 genes in BC pharmacogenomics.

## INTRODUCTION

Breast cancer (BC) is a heterogeneous disease characterized by an intricate interplay between different biological aspects such as ethnicity, genomic alterations, gene expression deregulation, hormone disruption, signaling pathway alterations and environmental determinants^1,2^. Over the last years, prevention, treatment and survival strategies have evolved favorably; however, there are BC profiles that remain incurable^3^. Nowadays, BC is the leading cause of cancer-related death among women (626,679; 15% cases) and the most commonly diagnosed cancer (2,088,849; 24% cases) worldwide^4^.

The development of large-scale DNA sequencing, gene expression, proteomics, large-scale RNA interference (RNAi) screens and large-scale CRISPR-Cas9 screens has allowed us to better understand the molecular landscape of oncogenesis. Significant progress has been made in discovering gene coding regions^5^, cancer driver genes^6,7^, cancer driver mutations^8,9^, germline variants^10^, driver fusion genes^11,12^, alternatively spliced transcripts^13^, expression-based stratification^14^, molecular subtyping^15^, biomarkers^16^, druggable enzymes^17^, cancer dependencies^18–21^, and drug sensitivity and resistance^22^.

Scientific advances made to date mark the era called the “end of the beginning” of cancer omics. In other words, each approach that was previously mentioned needs to be fully understood as a part of a complex network, analyzing the mechanistic interplay of signaling pathways, protein-protein interaction (PPi) networks, enrichment maps, gene ontology (GO), deep learning, molecular dependencies and genomic alterations per intrinsic molecular subtype: basal-like (estrogen receptor (ER)^-^, progesterone receptor (PR)^-^, human epidermal growth factor receptor 2 (Her2)^-^, cytokeratin 5/6^+^ and/or EGFR^+^); Her2-enriched (ER^-^, PR^-^, Her2^+^); luminal A (ER^+^ and/or PR^+^, Her2^-^, low Ki67); luminal B with Her2^-^ (ER^+^ and/or PR^+^, Her2^-^, low Ki67); luminal B with Her2^+^ (ER^+^ and/or PR^+^, Her2^-^, any Ki67); and normal like^23–29^. We will herein analyze previously prioritized genes/biomarkers by the Consensus Strategy (CS)^28^, the Pan-Cancer Atlas (PCA)^3,12,30–36^, the Pharmacogenomics Knowledgebase (PharmGKB)^37^ and the Cancer Genome Interpreter (CGI)^38^.

In our previous studies, López-Cortés *et al*. and Tejera *et al*., developed a Consensus Strategy that was proved to be highly efficient in the recognition of gene-disease association^28,39^. The main objective was to apply several bioinformatics methods to explore BC pathogenic genes. The CS identified both well-known pathogenic genes and prioritized genes that will be further explored through the OncoOmics approaches. On the other hand, The Cancer Genome Atlas (TCGA) has concluded the most sweeping cross-cancer analysis yet undertaken, namely the PCA project^31^. PCA reveals how genetic alterations, such as putative mutations, fusion genes, mRNA expression, copy number variants (CNVs) and protein expression collaborate in BC progression, providing insights to prioritize the development of new treatments and immunotherapies^3,12,30–36^. The CGI flags genomic biomarkers of drug response with different levels of clinical relevance^38^. Lastly, PharmGKB is a comprehensive resource that curates and spreads knowledge of the impact of clinical annotations on BC drug response^37,40^. PharmGKB collects the precise guidelines for the application of pharmacogenomics in clinical practice published by the European Society for Medical Oncology (ESMO), the National Comprehensive Cancer Network (NCCN), the Royal Dutch Association for the Advancement of Pharmacy (DPWG), the Canadian Pharmacogenomics Network for Drug Safety (CPNDS) and the Clinical Pharmacogenetics Implementation Consortium (CPIC)^41–44^. Hence, the aim of this study was to implement OncoOmics approaches to analyze genetic alterations, signaling pathways, PPi networks, protein expression, BC dependencies and enrichment maps in order to reveal essential genes/biomarkers to accelerate the development of precision medicine in BC.

## RESULTS

### OncoPrint of genetic alterations according to the Pan-Cancer Atlas

PCA has reported the clinical data of 1084 individuals with BC and it can be visualized in the Genomic Data Commons of the National Cancer Institute and in the cBioPortal^45,46^. In regard to molecular subtypes and tumor stages, 46% were lumina A, 18% luminal B, 7% Her2-enriched, 16% basal-like and 3% normal-like, whereas 17% were stage T1, 58% stage T2, 23% stage T3 and 2% stage T4 (Table S1).

Figure 1A shows the average frequency of genetic alterations per gene set. The average frequency of the PCA gene set was 1.3, followed by CS gene set (1.2), PharmGKB/CGI gene set (1.1), BC driver genes (0.8) and non-cancer genes (0.4) (Table S2). Significant p-values (p < 0.001) were found among all gene sets. Therefore, the fact that gene sets of interest (CS, PCA and PharmGKB/CGI) presented an average frequency of genetic alterations greater than the non-cancer gene set and the BC driver gene set indicates that we are analyzing potential essential genes in BC. Figure 1B shows the percentage of genetic alterations per type. The most common genetic alterations were mRNA upregulation (55.8%), CNV amplification (17.1%) and missense mutations (8.4%). Figure 1C shows the ratio of genetic alterations in the 230 genes per sample and molecular subtype. Basal-like had the highest ratio (n = 33), followed by Her2-enriched (29), luminal B (24), normal-like (17) and luminal A (15). The ratio of all BC samples was 19.6. Figure 1D shows the ratio of genetic alterations in the 230 genes per sample and tumor stage. Stage T2 had the highest ratio (23), followed by T3 (22), T1 (17) and T4 (8). Figures 1E and 1F show the percentage of genetic alterations per subtype and tumor stage, respectively. mRNA upregulation and CNV amplification were the most common alterations in all molecular subtypes and tumor stages.

**Figure 1.**
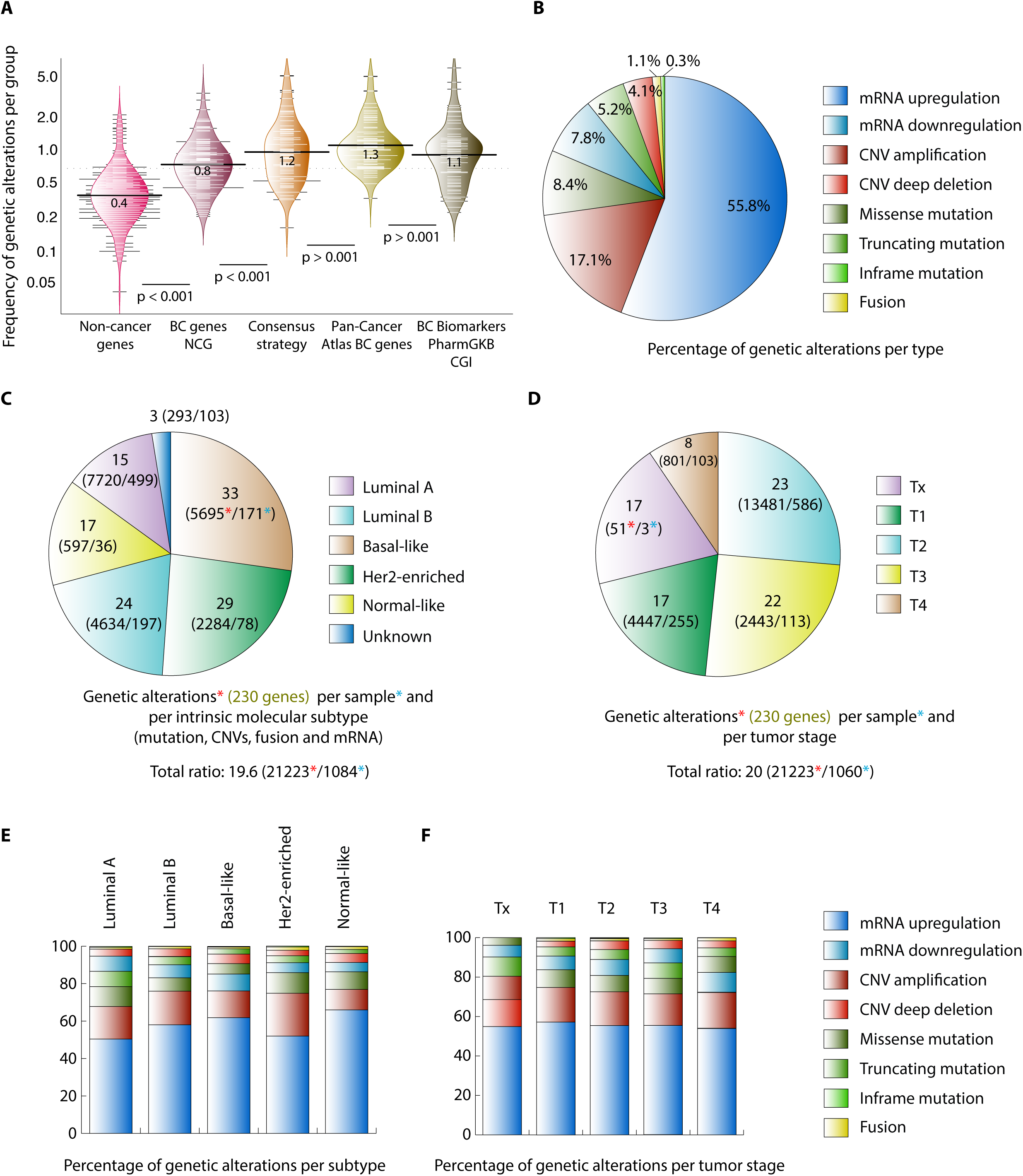
Genetic alterations of the breast cancer cohort according to PCA. (A) Frequency of genetic alterations per gene set (non-cancer genes, BC driver genes according to the Network of Cancer Genes, Consensus Strategy, BC genes according to PCA, BC biomarkers according to the PharmGKB and CGI). (B) Percentage of genetic alterations per type. (C) Ratio of genetic alterations per intrinsic molecular subtype. (D) Ratio of genetic alterations per tumor stage. (E) Percentage of genetic alterations per type and per molecular subtype. (F) Percentage of genetic alterations per type and per tumor stage.

Figure 2 shows the ranking of genes with the greatest number of genetic alterations per molecular subtype and tumor stage. Regarding molecular subtypes, *PIK3CA* was the most altered gene in luminal A, *CCND1* in luminal B, TP53 in basal-like and normal-like, and *ERBB2* in Her2-enriched, with significant p-values < 0.001 (Figure 2A). On the other hand, the most altered genes per tumor stage were *PIK3CA* in stage T1, TP53 in stages T2 and T3, and *ERBB2* in stage T4, with significant p-value < 0.001 (Figure 2B). Figures 2C, 2E, 2G, 2I and 2K show the top mutated genes, CNV amplified genes, CNV deep deleted genes, mRNA upregulated genes and mRNA downregulated genes per molecular subtype, respectively (Tables S3-S7). On the other hand, Figures 2D, 2F, 2H, 2J and 2L show the top mutated genes, CNV amplified genes, CNV deep deleted genes, mRNA upregulated genes and mRNA downregulated genes per tumor stage, respectively (Tables S8-S13).

**Figure 2.**
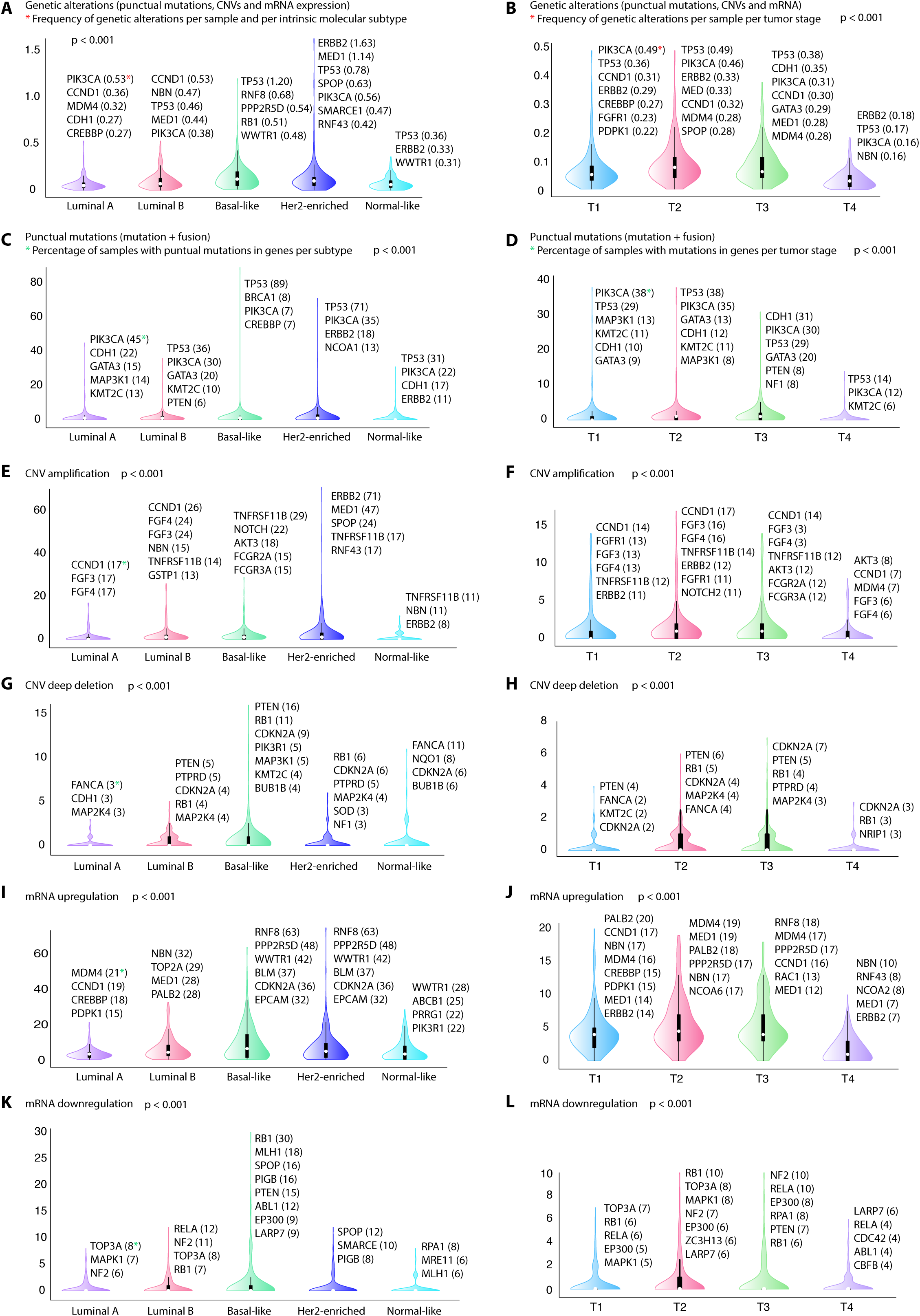
Ranking of genes with the highest number of genetic alterations per molecular subtype and tumor stage. (A) Frequency of genetic alterations (punctual mutations, copy number variants and mRNA expression) per molecular subtype. (B) Frequency of genetic alterations per tumor stage. (C) Frequency of punctual mutations per molecular subtype. (D) Frequency of punctual mutations per tumor stage. (E) Frequency of CNV amplifications per molecular subtype. (F) Frequency of CNV amplifications per tumor stage. (G) Frequency of CNV deep deletions per molecular subtype. (H) Frequency of CNV deep deletions per tumor stage. (I) Frequency of mRNA upregulation per molecular subtype. (J) Frequency of mRNA upregulation per tumor stage. (K) Frequency of mRNA downregulation per molecular subtype. (L) Frequency of mRNA downregulation per tumor stage.

Regarding the first OncoOmics approach, Figure 3A shows an OncoPrint of 73 genes with a number of genetic alterations greater than the average (> 86). For this analysis driver mutations were taken into account, discarding passenger mutations (Figure S1 and Table S14). Figure 3B shows a circos plot of interactions between molecular subtypes and genetic alterations of the 73 most altered genes. mRNA downregulated plus CNV deep deleted genes and mRNA upregulated plus CNV amplified genes were more related with basal-like, whereas fusion genes, and driver mutations were more related with Her2-enriched. Finally, Figure 3C shows a circos plot of interactions between tumor stages and genetic alterations of the 73 most altered genes. Fusion genes, mRNA downregulated plus CNV deep deleted genes, and mRNA upregulated plus CNV amplified genes were more related with stage T4, whereas driver mutations were more related with stage T3.

**Figure 3.**
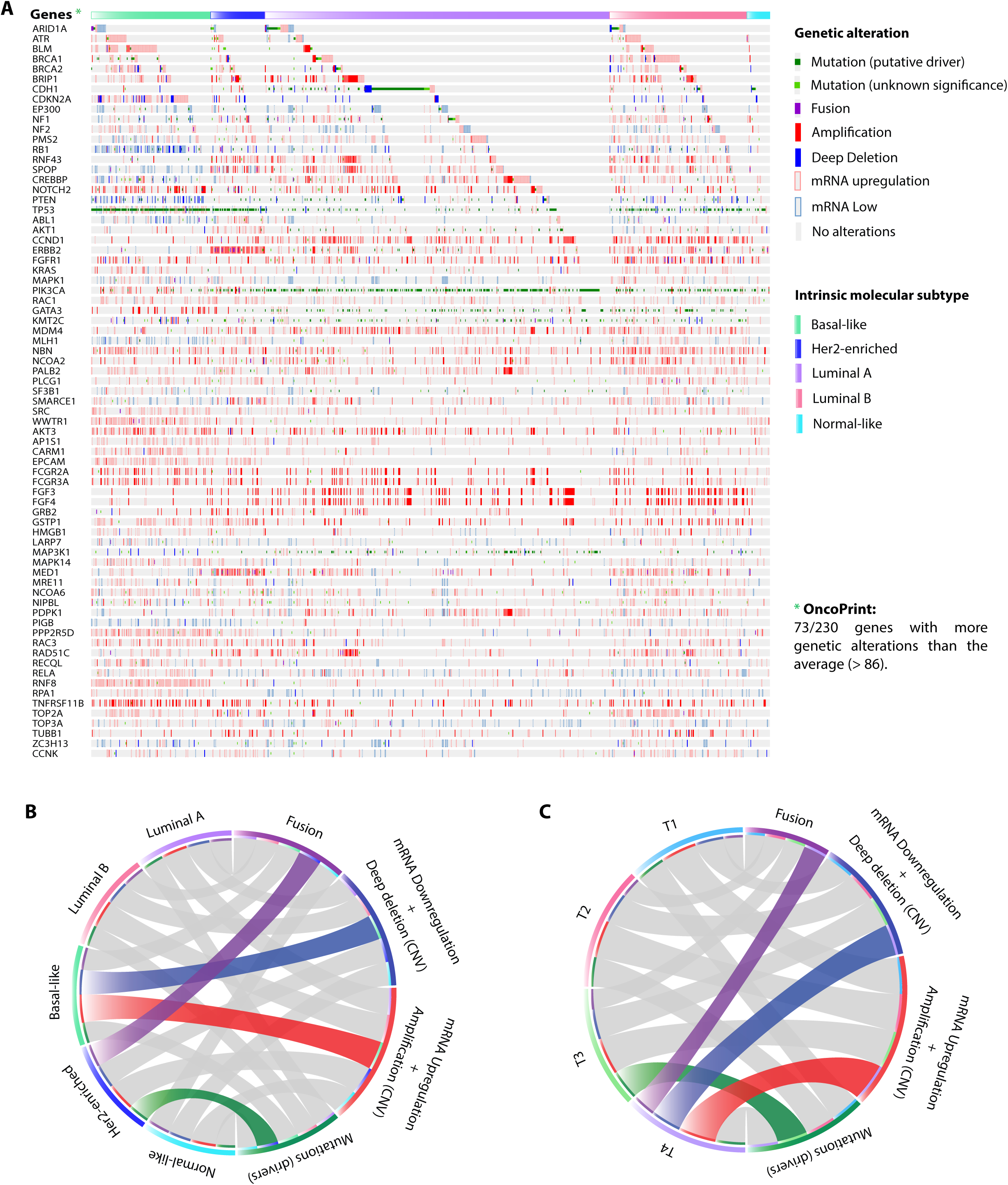
OncoPrint of genetic alterations according to the Pan-Cancer Atlas. (A) OncoPrint of genes with more genetic alterations than the average (>86) per molecular subtype. (B) Circos plot between molecular subtypes and the highest number of genetic alterations (fusion genes, mRNA downregulation plus CNV deep deletion, mRNA upregulation plus CNV amplification and driver mutations). (C) Circos plot between tumor stages and the highest number of genetic alterations.

### Pathway enrichment analysis

The pathway enrichment analysis was performed using David Bioinformatics Resource to obtain integrated information from the Kyoto Encyclopedia of Genes and Genomes (KEGG)^47–50^. The enrichment analysis of signaling pathways was carried on in the 230 genes, obtaining more than 50 terms with a false discovery rate (FDR) < 0.01 (Table S15). Subsequently, genetic alterations of genes that make up each signaling pathway were analyzed according to the molecular subtype and tumor stage. Figure 4A shows a circos plot correlating molecular subtypes with signaling pathways (Table S16). NF-kappa ß, NOD-like receptor, adipocytokine, GnRH, RIG-like receptor, TNF, TGFß, FOXO, glucagon, MAPK, prolactin, cAMP, PI3K-AKT, neurotrophin, VEGF, notch, p53, sphingolipid and Wnt signaling pathways were more altered in basal-like; estrogen, HIF1, toll-like receptor, ras, insulin, T-cell receptor, rap1, ERBB, AMPK, chemokine, B-cell receptor, mTOR, Fc-epsilon RI, Jak-STAT, phosphatidylinositol and thyroid hormone signaling pathways were more altered in Her2-enriched; and Hippo signaling pathway in normal-like. On the other hand, Figure 4B shows the ranking of the most altered signaling pathways per molecular subtype. Jak-STAT signaling pathway was more altered in luminal A; Wnt signaling pathway in luminal B; p53 signaling pathway in basal-like; ERBB signaling pathway in Her2-enriched; and Hippo signaling pathway in normal-like (Table S17).

**Figure 4.**
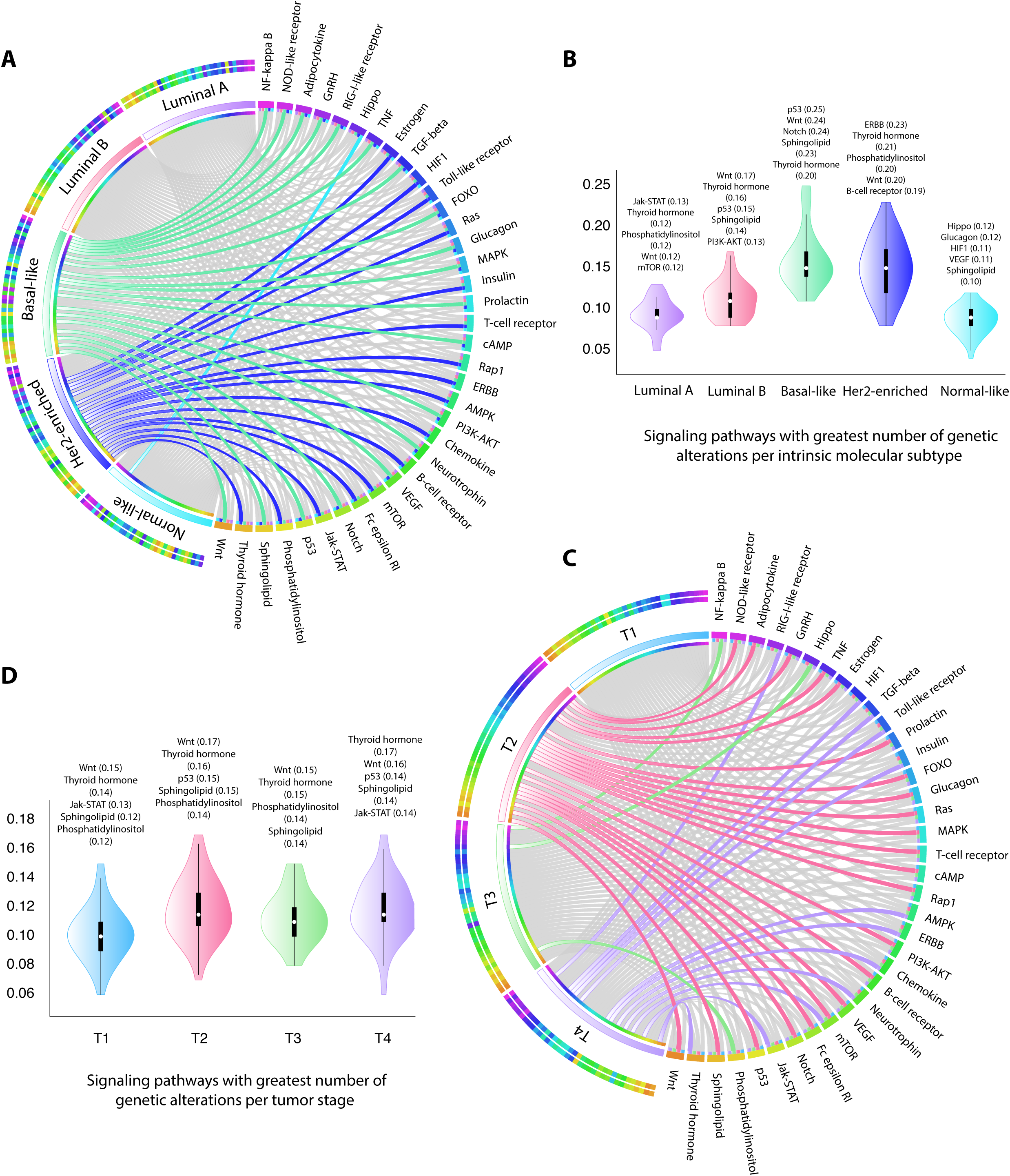
Pathway enrichment analysis per molecular subtype and tumor stage. (A) Circos plot between molecular subtypes and the most altered genetic pathways. (B) Violin plots showing the frequency of the most altered signaling pathways per molecular subtype. (C) Circos plot between tumor stages and the most altered genetic pathways. (D) Violin plots showing the frequency of the most altered signaling pathways per tumor stage.

Figure 4C shows a circos plot correlating tumor stages with signaling pathways according to the frequency of genetic alterations (Table S16). NOD-like receptor, adipocytokine, GnRH, TNF, estrogen, prolactin, FOXO, glucagon, ras, MAPK, T-cell receptor, cAMP, rap1, PI3K-AKT, B-cell receptor, VEGF, mTOR, Fc epsilon RI, NOTCH, p53, sphingolipid and Wnt signaling pathways were more altered in stage T2; NF-kappa ß, Hippo and phosphatidylinositol signaling pathways were more altered in stage T3; and RIG-like receptor, HIF1, TGFß, toll-like receptor, insulin, AMPK, ERBB, chemokine, neurotrophin, mTOR, jak-STAT and thyroid hormone signaling pathways were more altered in stage T4. On the other hand, Figure 4D shows the ranking of the most altered signaling pathways per tumor stage. Wnt signaling pathway was more altered in stages T1, T2 and T3; and thyroid hormone signaling pathway was more altered in stage T4 (Table S18).

### Protein-protein interaction network

Regarding the second OncoOmics approach, the PPi network was performed to better understand BC behavior using the String Database and Cytoscape^51,52^. With the indicated cutoff of 0.9, the final interaction network had 258 nodes conformed by 198 (86%) genes from the CS, PCA and PharmGKB/CGI gene sets, and enriched with 60 previously known BC driver genes. Regarding the OncoPrint genes, 65 (89%) nodes integrated this network (Figure 5A). On the other hand, out of the 258 genes that make up our String PPi network, 16 (6%) genes and 18 edges were part of the OncoPPi BC network^53,54^. The degree centrality made it possible to establish a significant correlation (Spearman p < 0.05) between our String PPi network and the OncoPPi BC network (Figure 5B).

**Figure 5.**
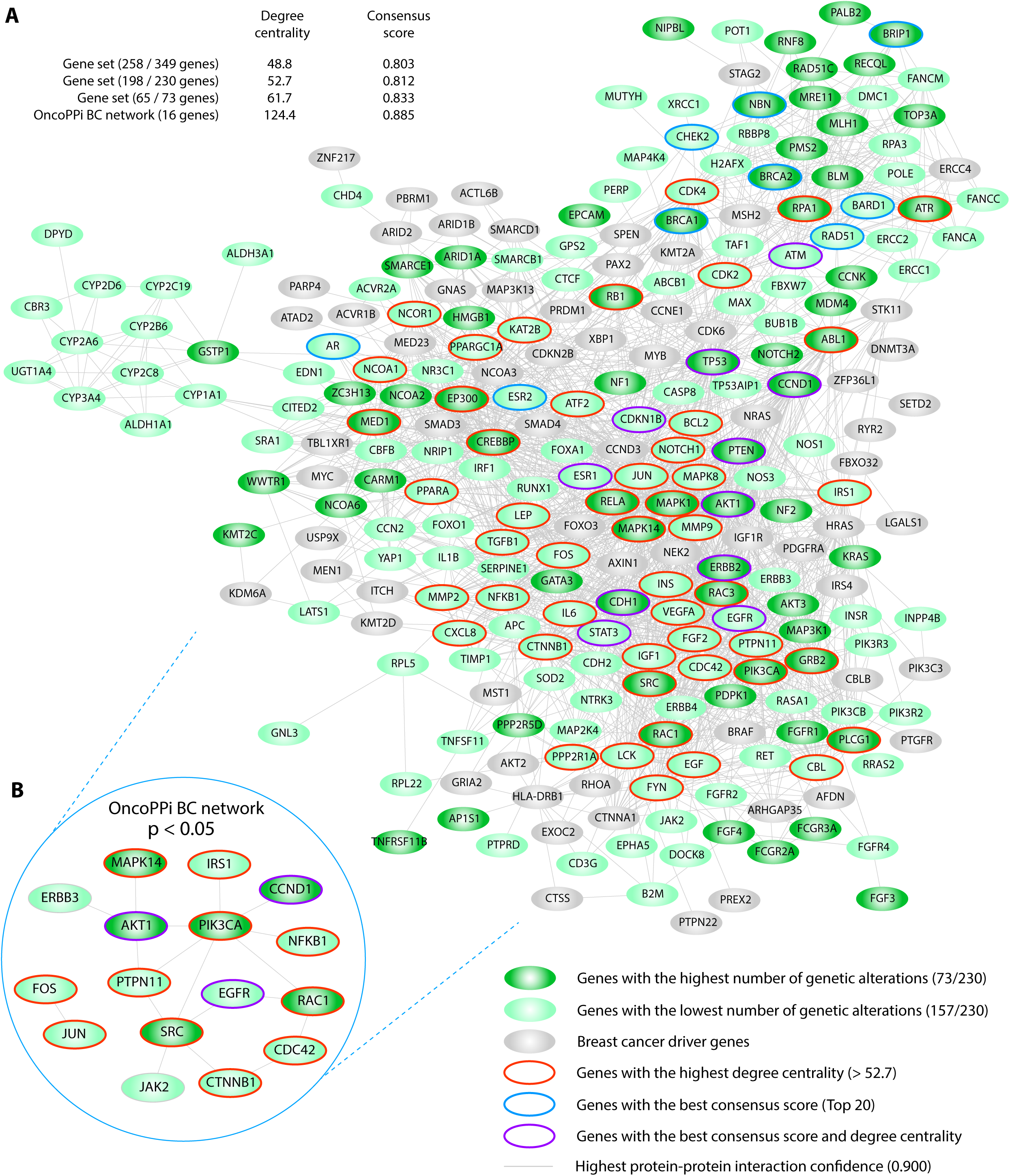
Breast cancer integrated network. (A) Network composed of BC driver genes and genes of our study (PCA gene set, consensus strategy gene set and PharmGKB gene set. (B) Significant correlation (p < 0.05) of degree centrality and consensus score between the OncoPPi BC network and or BC integrated network.

Considering the degree centrality and the consensus score of our previous study^28^, there was enrichment among sub-networks (Figures 5A and 5B). The average of degree centrality of the 258 nodes network was 48.8; out of the 198 nodes network was 52.7; out of the 65 nodes network was 61.7; and out of the OncoPPi BC network was 124.4. Meanwhile, the average of consensus score of the 258 nodes network was 0.803, out of the 198 nodes network was 0.812, out of the 65 nodes network was 0.833, and out of the OncoPPi BC network was 0.885. Additionally, the second OncoOmics approach was made up of genes with the highest degree centrality (> 52.7) such as *TP53*, *AKT1*, *SRC*, *CREBBP*, *EP300*, *JUN*, *CTNNB1*, *PIK3CA*, *RAC1* and *EGFR*, genes with the highest consensus score such as *TP53*, *ESR1*, *CCND1*, *BRCA2*, *BRCA1*, *ERBB2*, *CHEK2*, *AR*, *MYC* and *PTEN*, and genes with both of them such as *TP53*, *ESR1*, *CCND1*, *ERBB2*, *PTEN*, *CDKN1B*, *ATM*, *AKT1*, *STAT3*, *CDH1* and *EGFR* (Table S19).

### Protein expression analysis

The third OncoOmics approach was related to the expression analysis of the 230 proteins. Figure 6A shows 43 proteins with significant high expression (Z-scores ≥2) and low expression (Z-scores ≤-2) analyzed with the reverse-phase protein array (RPPA) and mass spectrometry, according to TCGA. The top ten proteins with the highest expression levels in a cohort of 994 individuals were *ERBB2*, *SERPINE2*, *CDH2*, *CCND1*, *EGFR*, *ERCC1*, *IRS1*, *NOTCH1*, *ERBB3* and *INPP4B*, and the ones with the lowest expression levels were *CDH1*, *ATM*, *JAK2*, *MAPK1*, *AKT1*, *AKT3*, *MAPK14*, *ABL1*, *CTNNB1* and *IRF1* (Table S20). On the other hand, the Human Protein Atlas (HPA) presented a map of the human tissue proteome based on tissue microarray-based immunohistochemistry. HPA has analyzed 202 (88%) of the 230 proteins of our study, classifying the protein expression in high, medium, low and non-detected. As a result, *RAC1*, *GJB2*, *MED1*, *PIK3CA*, *PIK3R3*, *FGFR2*, *HCFC2*, *MAP2K4*, *NQO2* and *RAC3* were proteins with high and medium expression in normal tissue, and low and non-detected expression in BC tissue, acting as tumor suppressor genes. Meanwhile, *CDK2*, *CYP2D6*, *NCOR1*, *RRM1*, *FOXA1* and *TOP2A* were proteins with high and medium expressions in BC tissue, and low and non-detected expressions in normal tissue, acting as oncogenes (Figure 6B and Table S21). Lastly, according to the HPA, Figure 6C shows the overall survival analysis of *RAD51*, *PERP* and *MORC4* as BC biomarkers with unfavourable prognosis and p < 0.001 (Table S22)^55,56^. All these altered proteins made up the third OncoOmics approach.

**Figure 6.**
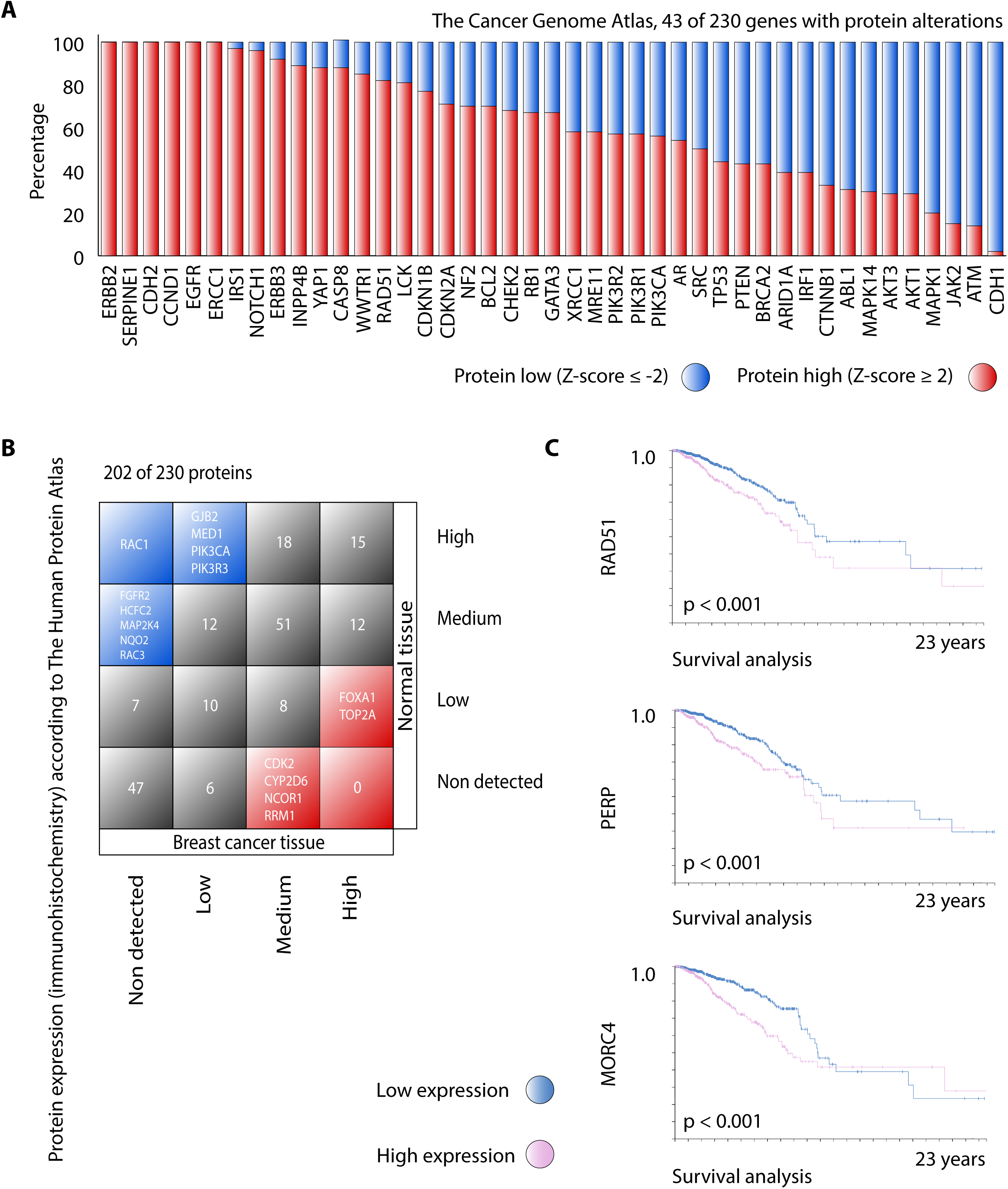
Analysis of protein expression. (A) Ranking of genes with the highest number of protein alterations (high and low expression with Z-score ≥ 2) according to The Cancer Genome Atlas. (B) Comparison of protein expression levels between BC tissue and normal tissue according to The Human Protein Atlas. (C) Overall survival of genes with prognosis unfavorable (p < 0.001) in BC according to The Human Protein Atlas.

### Breast cancer dependency map

The fourth OncoOmics approach consisted in identifying genes that are essential for cancer cell proliferation and survival performing systematic loss-of-function screens in a large number of well-annotated cancer cell lines and BC cell lines representing the tumor heterogeneity^18–21^. Figure 7A shows the distribution of dependency scores of 227/230 genes through DEMETER2, an analytical framework for analyzing genome-scale RNAi loss-of-function screens in 73 BC cell lines (Table S23). Our results showed 563 dependencies with at least one score _≤_ −1 in 57 (25%) essential genes. The top 10 genes with the greatest number of significant dependency scores in BC cell lines were *RPL5* (68; 93%), *SF3B1* (67; 92%), *RPA1* (61; 84%), *RRM1* (53; 73%), *BUB1B* (26; 36%), *RPA3* (25; 34%), *RAD51* (23; 32%), *PPP2R1A* (21; 29%), *CHD4* (19; 26%) and *POLE* (13, 18%). At the same time, Figure 7A shows the distribution of dependency scores of 217/230 genes through CERES, an analytical framework for analyzing genome-scale CRISPR-Cas9 loss-of-function screens in 28 BC cell lines (Table S24). Our results showed 310 dependencies with at least one score _≤_ −1 in 34 (16%) essential genes. The top 10 genes with the greatest number of significant dependency score in BC cell lines were *RPA1* (27; 96%), *RRM1* (27; 96%), *TOP2A* (26; 93%), *BUB1B* (24; 86%) *CTCF* (24; 86%), *POLE* (23; 82%), *SF3B1* (19; 68%), *RPL5* (17; 61%), *CCND1* (13; 46%) and *SOD2* (13; 46%). Figure 7B shows the distribution of dependency scores of DEMETER2 and CERES per molecular subtype. The genome-scale RNAi loss-of-function screens detected 165 (29%) dependencies in 19 Her2-enriched cell lines (ratio = 8.7), 110 (20%) in 13 luminal A cell lines (8.5), 57 (10%) in 7 luminal B cell lines (8.1), and 231 (41%) in 34 basal-like cell lines (6.8), whereas the genome-scale CRISPR-Cas9 loss-of-function screens detected 85 (27%) dependencies in 7 luminal A cell lines (ratio = 12.1), 176 (15%) in 16 basal-like cell lines (11), and 49 (16%) in 5 Her2-enriched cell lines (9.8). Figure 7C shows violin plots of dependencies per molecular subtype. DEMETER2 has detected a greatest number of significant dependencies in basal-like, followed by Her2-enriched, luminal A and luminal B, whereas CERES has detected a greatest number of significant dependencies in basal-like, followed by luminal A and Her2-enriched. Figure 7D shows a Venn diagram of 66 essential genes with at least one significant dependency in different molecular subtypes, where 22 were strongly selective genes, 26 were common essential genes, and 5 were both of them in all cancer cell lines (Figure 7E).

**Figure 7.**
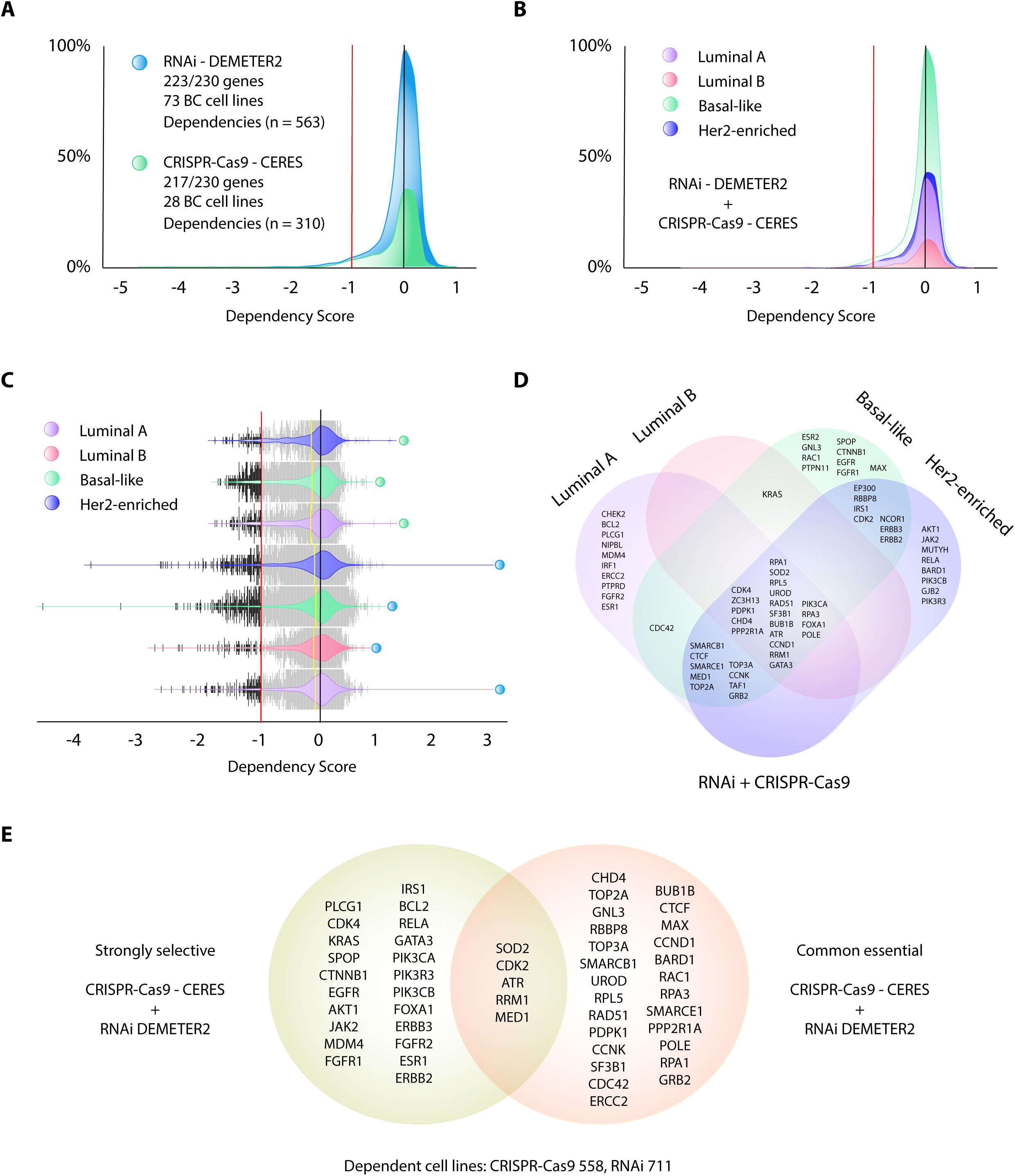
Analysis of dependencies in BC cell lines. (A) Dependency score of BC gene sets using RNAi DIMETER2 and CRISPR-Cas9 CERES algorithms in BC cell lines. (B) Dependency score of BC gene sets per molecular subtypes. (C) Violin plots of dependencies per molecular subtypes. All significant dependencies < −1 are in black. (D) Venn diagram of genes with at least one dependency < −1 in cell lines belonging to each molecular subtype. (E) Venn diagram of strongly selective and common essential genes in all cancer cell lines.

### OncoOmics approaches to reveal essential genes in BC

Figure 8A shows a Venn diagram integrated by the OncoOmics essential genes, the most relevant genes of the CS, PCA and PharmGKB/CGI gene sets per approach. *RAC1*, *AKT1*, *CCND1*, *PIK3CA* and *ERBB2* were relevant genes in all OncoOmics approaches; *CDH1*, *MAPK14*, *TP53*, *MAPK1*, *SRC* and *RAC3* were relevant genes in the OncoPrint, networking and protein expression analyses; *PLCG1* and *GJB2* were relevant genes in the OncoPrint, networking and DepMap analyses; *MED1*, *TOP2A* and *GATA3* were relevant genes in the DepMap, OncoPrint and protein expression analyses; *BCL2*, *CTNNB1*, *EGFR* and *CDK2* were relevant in the DepMap, networking and protein expression analyses; *EP300* and *CREBBP* were relevant in the networking and the OncoPrint analyses; *PTEN*, *MRE11*, *CDKN2A*, *WWNTR1*, *ABL1*, *BRCA2*, *NF2*, *AKT3*, *ARDID1A* and *RB1* were relevant in the OncoPrint and protein expression analyses; *RPA1*, *TOP3A*, *FGFR1*, *SF3B1*, *ATR*, *KRAS*, *PDPK1*, *RELA*, *SMARCE1*, *SPOP*, *CCNK* and *MDM4* were relevant in the DepMap and OncoPrint analyses; *CDKN1B*, *LCK* and *NOTCH1* were relevant in the networking and protein expression analyses, *CDK4* and *ESR1* were relevant in the DepMap and networking analyses; and *RAD51*, *IRS1*, *FGFR2*, *JAK2*, *RRM1*, *PIK3R3*, *FOXA1* and *ERBB3* were relevant in the DepMap and protein expression analyses (Table S25).

**Figure 8.**
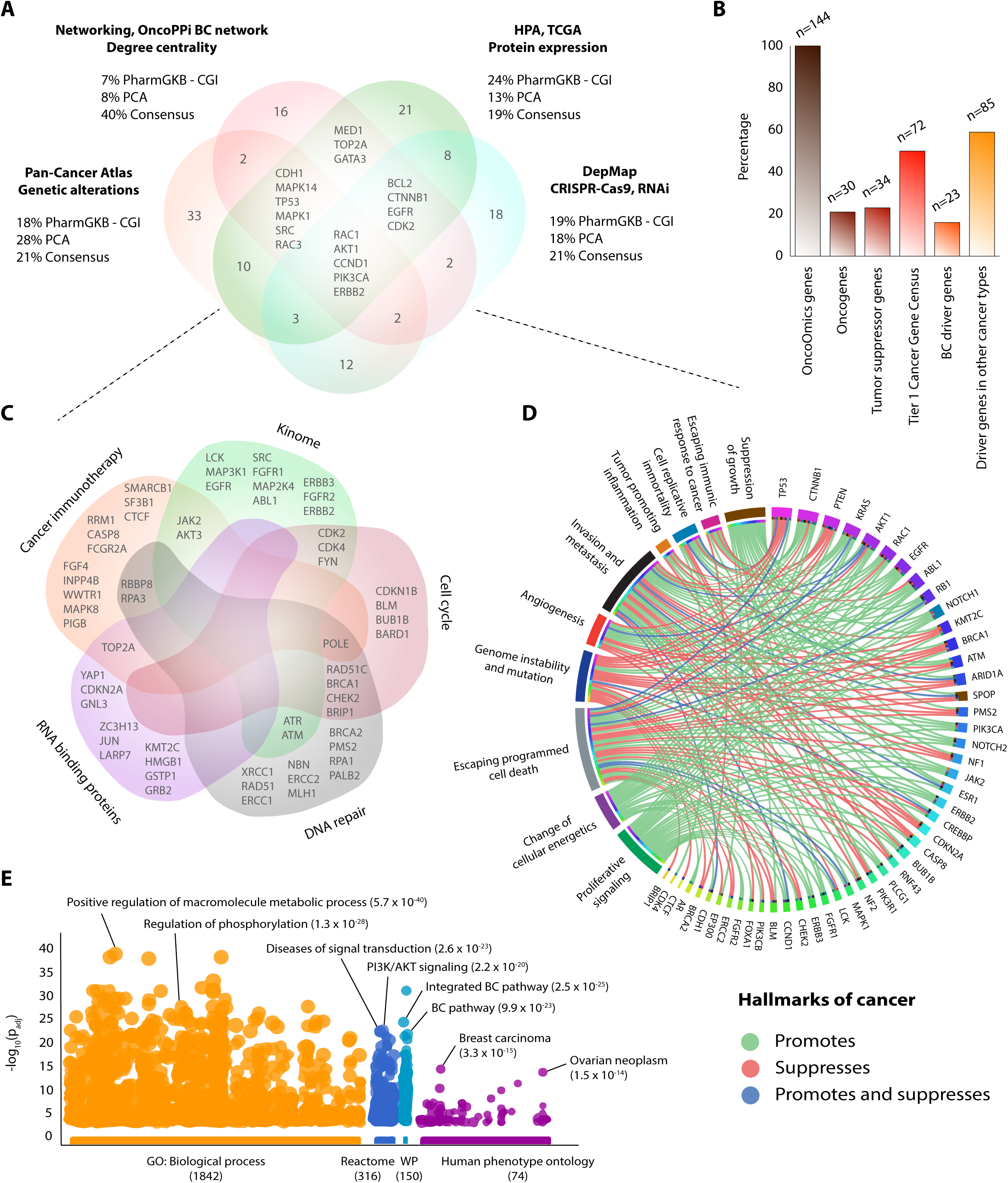
The OncoOmics essential genes of breast cancer. (A) Venn diagram of the most relevant genes per genomics approach (PCA genetic alterations, networking, protein expression and DepMap). (B) Percentage of oncogenes, tumor suppressor genes, tier 1 genes, BC driver genes and driver genes in other cancer types. (C) Venn diagram of the most relevant genes related with cancer immunotherapy, kinome, cell cycle, DNA repair and RNA-binding proteins. (D) Circos plot of the hallmarks of cancer genes. (E) Most significant g:Profiler features of the most relevant genes according to the gene ontology biological processes, Reactome pathways, wikipathways and the human phenotype ontology.

Out of the 144 OncoOmics essential genes, 21% were oncogenes, 24% were tumor suppressor genes, 50% were tier 1, according to the Cancer Gene Census (COSMIC)^60^, and 59% were driver genes in other types of cancer, according to The Network of Cancer Genes^61^ (Figure 8B). On the other hand, *FGF4*, *INPP4B*, *WWNTR1*, *MAPK8*, *PIGB*, *RRM1*, *CASP8*, *FCGR2A*, *SMARCB1*, *SF3B1* and *CTCF* were cancer immunotherapy genes^62^; *LCK*, *MAP3K1*, *EGFR*, *SRC*, *FGFR1*, *MAP2K4*, *ABL1*, *ERBB3*, *FGFR2* and *ERBB2* were kinome genes^63^; *CDKN1B*, *BLM*, *BUB1B* and *BARD1* were cell cycle genes^64^; *XRCC1*, *RAD51*, *ERCC1*, *NBN*, *ERCC2*, *MLH1*, *BRCA2*, *PMS2*, *RPA1* and *PALB2* were DNA repair genes^65^; lastly, *YAP1*, *CDKN2A*, *GNL3*, *ZC3H13*, *JUN*, *LARP7*, *KMT2C*, *HMGB1*, *GSTP1* and *GRB2* were RNA-binding proteins (RBPs) (Figure 8C and Table S26)^66^.

Figure 8D shows a circos plot of the 48 (33%) OncoOmics essential genes that are hallmarks of cancer. The top 10 genes with the greatest number of interactions with the hallmarks of cancer were *TP53*, *CTNNB1*, *PTEN*, *KRAS*, *AKT1*, *RAC1*, *EGFR*, *ABL1*, *RB1* and *NOTCH1*. Suppression of growth was promoted by *AKT1*, *CTNNB1*, *PTEN*, *RB1* and *TP53*; escaping immune response to cancer was promoted by *CTNNB1*, *EGFR* and *RAC1*, and suppressed by *ABLI*, *PTEN* and *TP53*; cell replicative immortality was promoted by *CTNNB1*, *KRAS* and *NOTCH1*, suppressed by *PTEN*, and promoted/suppressed by *TP53*; tumor promoting inflammation was promoted by *KRAS* and suppressed by *TP53*; metastasis was promoted by *ABL1*, *CTNNB1*, *EGFR*, *KRAS*, *RAC1* and *RB1*, suppressed by *PTEN* and *TP53*, and promoted/suppressed by *AKT1*; angiogenesis was promoted by *ABL1*, *CTNNB1*, *EGFR*, *KRAS*, *NOTCH1* and *RAC1*, suppressed by *TP53* and promoted/suppressed by *AKT1*; genome instability was promoted by *ABL1* and *RB1*, and suppressed by *AKT1*, *CTNNB1*, *PTEN*, *RAC1* and *TP53*; escaping programmed cell death was promoted by *AKT1*, *CTNNB1*, *EGFR*, *NOTCH1* and *RAC1*, suppressed by *PTEN*, and promoted/suppressed by *KRAS*, *RB1* and *TP53*; change of cellular energetics was promoted by *ABL1*, *AKT1*, *CTNNB1*, *EGFR*, *KRAS*, *NOTCH1*, *PTEN*, *RB1* and *TP53*; finally, proliferative signaling was promoted by *ABL1*, *AKT1*, *CTNNB1*, *EGFR*, *KRAS*, *NOTCH* and *RAC1* (Table S27).

### Enrichment map of the OncoOmics essential genes in BC

Figure 8E shows the enrichment map of the 144 OncoOmics essential genes in BC. g:Profiler searches for a collection of gene sets representing pathways, networks, GO terms and disease phenotypes^67^. The most significant GO: biological process with a FDR < 0.001 was positive regulation of macromolecule metabolic process (Table S28); the most significant GO: molecular function was phosphatidylinositol 3-kinase activity (Table S29); the most significant Reactome pathway was generic transcriptor pathway (Table S30)^68^; additionally, the most significant disease, according the Human Phenotype Ontology, was breast carcinoma (Table S31)^69^. Subsequently, g:Profiler annotations were analyzed with the EnrichmentMap software and visualized using Cytoscape, in order to generate network interactions of the most relevant GO: biological processes (Figure S2) and Reactome pathways (Figure 9) related to immune system, tyrosine kinase, cell cycle and DNA repair pathways^52,67^.

**Figure 9.**
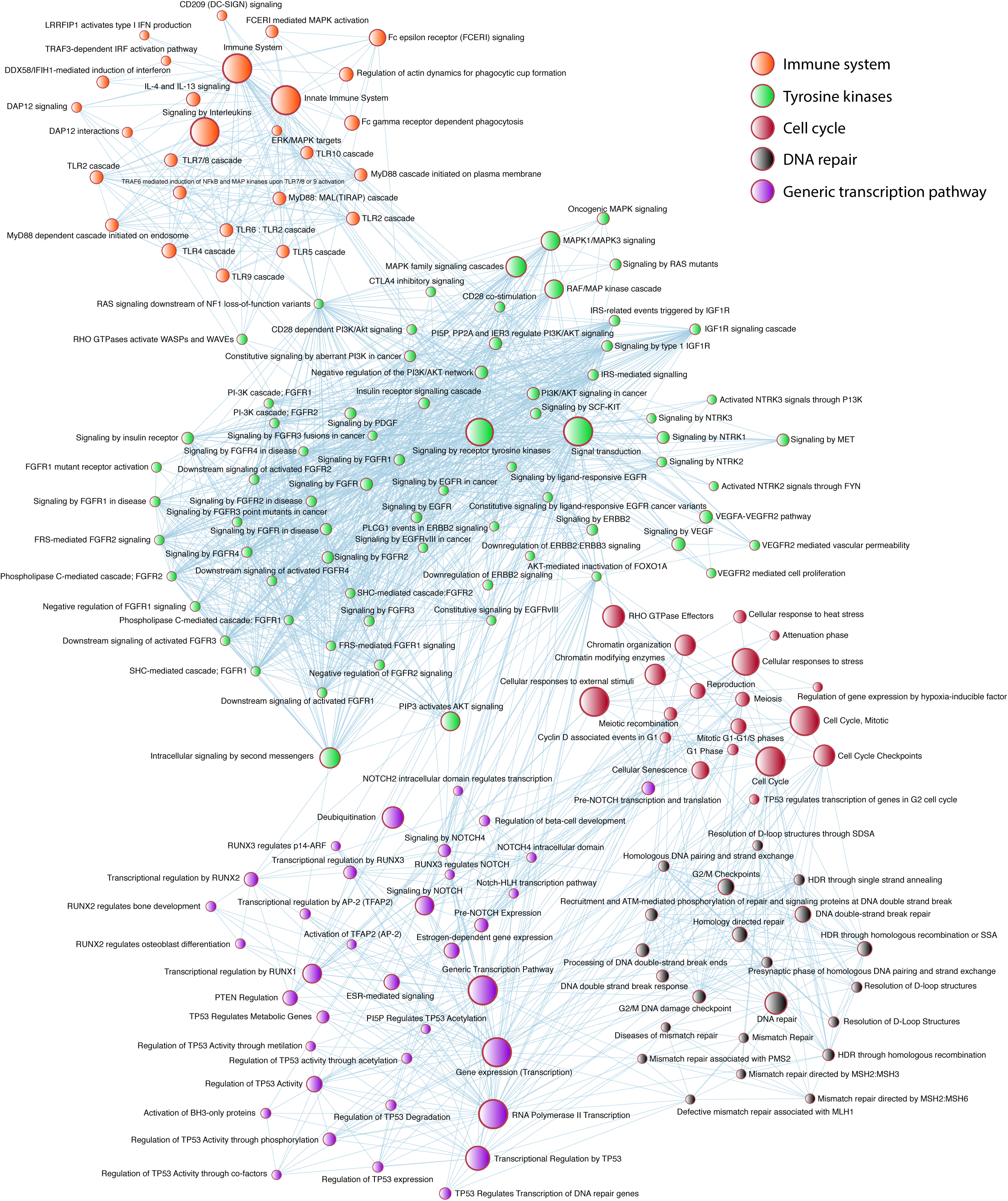
Pathway enrichment analysis of the most relevant genes using g:Profiler and EnrichmentMap. Most significant Reactome pathways related to immune system, tyrosine kinases, cell cycle, DNA repair and genetic transcription.

### Precision medicine

Figure 10 shows the current status of clinical trials for BC, according to the Open Targets Platform^70^. There are 111 drugs that are being analyzed in 3151 clinical trials in 39/230 genes. The top 10 genes with the highest number of clinical trials in process or completed were *TUBB1*, *ERBB2*, *ESR1*, *TOP2A*, *EGFR*, *ESR2*, *VEGFA*, *CDK4*, *POLE* and *RRM1*. The greatest number of clinical trials was in phase 2. Small molecules were the most analyzed type of drug, followed by antibodies and proteins. Lastly, the target classes with the greatest number of clinical trials were tyrosine kinases, structural proteins and nuclear hormone receptors (Table S32).

**Figure 10.**
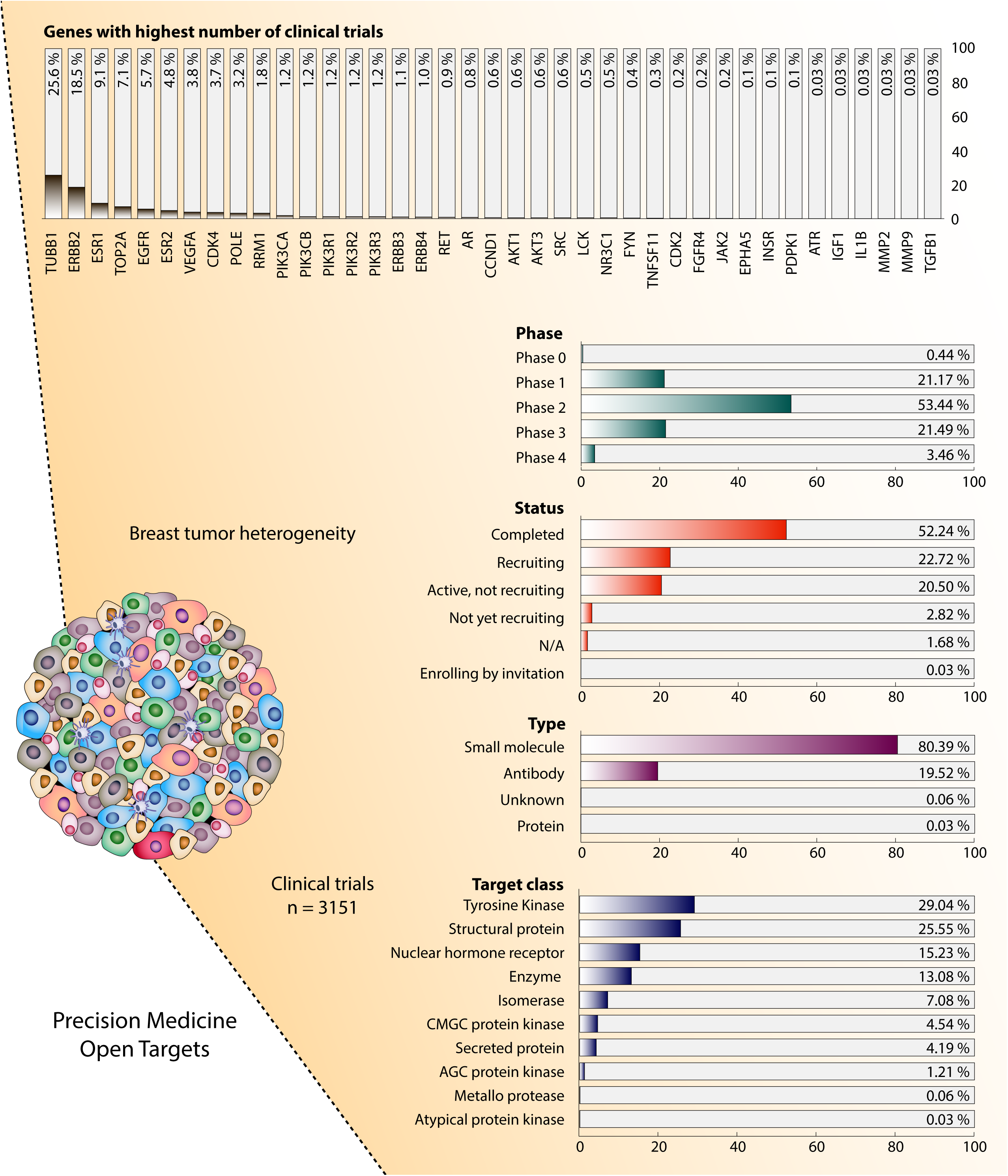
A panoramic view of clinical trial features in breast cancer.

Regarding precise guidelines for the application of BC pharmacogenomics in clinical practice, PharmGKB details 154 clinical annotations in 70/230 (30%) genes (Table S33)^41–44^; the CGI details 76 clinical annotations in 26/230 (11%) genes (Table S34)^71^; and PCA details 648 clinical annotations in 14/230 (6%) genes (Table S35)^72^.

Additionally, Figure S3 shows a drug-gene interaction matrix conformed by 109 clinical annotations in phase 4, according to the OTP; 9 clinical annotations in levels 1A, 2A and 2B, according to PharmGKB; 9 clinical annotations approved by the US Food and Drug Administration (FDA), according to CGI; and 648 clinical annotations, according to PCA.

## DISCUSSION

In this study we proposed a compendium of OncoOmics approaches that analyze genetic alterations, protein expression, signaling pathways, PPi networks, enrichment maps, gene ontology and dependency maps in three gene sets. The first gene set was taken from our previous study where we developed a Consensus Strategy that was proved to be highly efficient in the recognition of BC pathogenic genes^28^. The second gene set was taken from several studies of PCA, which provides a panoramic view of the oncogenic processes that contributes to BC progression^3,12,30–36^. The third gene set was taken from the CGI and PharmGKB. On the one hand, the CGI flags genomic biomarkers of drug response with different levels of clinical relevance^38^. On the other hand, PharmGKB collects clinical annotations applied in BC patients and taken from the NCCN, ESMO, CPNDS, DPWG and CPIC guidelines^41–44^. Finally, the compendium of these 230 potential essential genes in BC was analyzed through four different OncoOmics approaches.

The first OncoOmics approach consisted in the analysis of genetic alterations using the PCA data^45,46^. The frequency of genetic alterations in the CS (average = 1.2), PCA (1.3) and PharmGKB/CGI (1.1) gene sets were higher than the non-cancer gene set (0.4) and the previously known BC driver genes (0.8). This means that these 230 genes had a greater number of genetic alterations and might be strongly associated with BC (Figure 1A). The most common genetic alterations in a cohort of 994 individuals were mRNA upregulation, CNV amplification and missense mutations. Molecular subtypes with the greatest number of genetic alterations were basal-like, Her2-enriched, luminal B, normal-like and luminal A, whereas tumor stages with the greatest number of genetic alterations were T2, T3, T1 and T4 (Figures 1B-F). Genes with the greatest number of genetic alterations per subtype were *PIK3CA* in luminal A, *CCND1* in luminal B, *TP53* in basal-like and normal-like, and *ERBB2* in Her2-enriched (Figure 2A), whereas *PIK3CA* was the most altered gene in stage T1, *TP53* in stages T2 and T3, and *ERBB2* in stage T4 (Figure 2B).

After a thorough analysis of genetic alterations in the 230 genes, the first OncoOmics approach was generated by an OncoPrint conformed by the top 73 genes with the greatest number of genetic alterations and with a frequency of alterations greater than the average (> 86) (Figure 3A). The top 10 most altered genes were *PIK3CA*, *TP53*, *MDM4*, *CCND1*, *NBN*, *MED1*, *CREBBP*, *PALB2*, *ERBB2* and *SPOP*^3,12,30–36^.

Subsequently, the enrichment analysis of signaling pathways was carried on taking into account all genetic alterations in the 230 genes using David Bioinformatics Resource and KEGG^47,50^. The signaling pathways with the greatest number of genetic alterations per intrinsic molecular subtype were Jak-STAT in luminal A, Wnt in luminal B, p53 in basal-like, ERBB in Her2-enriched and Hippo in normal-like (Figure 4B); and per tumor stage were Wnt in stages T1, T2 and T3, and thyroid hormone in stage T4 (Figure 4D).

Regarding the previously mentioned signaling pathways, Jak-STAT is involved in the control of processes, such as stem cell maintenance, hematopoiesis and inflammatory response. However, the mechanism underlying inappropriate Jak-STAT pathway activation is not well-known in BC^73^. The Wnt signaling pathway actively functions in embryonic development and helps in homeostasis in mature tissues by regulating cell survival, migration, proliferation and polarity^74^. The p53 tumor suppressor is the most frequently mutated gene in human cancer^75^, and acting as a transcription factor, the p53 signaling pathway plays a critical role in growth-inhibition, apoptosis, cell migration and angiogenesis^76^. The ERBB signaling pathway members form cell-surface receptors with extracellular domains yielding ligand-binding specificity^77^. Downstream signaling proceeds via tyrosine phosphorylation mediating signal transduction events that control cell survival, migration and proliferation. However, aberrant ERBB activation can increase transcriptional expression^78^. The Hippo pathway plays important roles in immune response, stem cell function and tumor suppression. However, alterations in this pathway are involved in the BC tumorigenesis and metastasis^79^. Lastly, the thyroid hormone signaling pathway is an important regulator of growth and metabolism. Nevertheless, deregulation of the T3 hormone levels could promote abnormal responsiveness of mammary epithelial cells developing BC^80^.

The second OncoOmics approach consisted in the PPi network analysis and its validation with the OncoPPi BC network. According to Li *et al*. and Ivanov *et al*.^54,81^, PPi with therapeutic significance can be revealed by the integration of cancer genes into networks. PPi regulates essential oncogenic signals to cell proliferation and survival, and thus, represents potential targets for drug development and drug discovery. Regarding our networking analysis, the final interaction network consisted in 258 nodes with an average of degree centrality of 48.8 and an average of consensus scoring of 0.803^28^; the sub-network integrated by 198 of 230 nodes had 52.7 of degree centrality and 0.812 of consensus scoring; finally, the sub-network integrated by 65 of 73 genes with the greatest number of genetic alterations had 61.7 of degree centrality and 0.833 of consensus scoring. Hence, a sub-network of genes with greatest number of genetic alterations presented a greater degree centrality and consensus scoring, suggesting that there is strong correlation between these genes and BC. Additionally, the oncogenomics validation showed a significant correlation between our String PPi network (Figure 5A) and the OncoPPi BC network (Figure 5B), identifying 16 nodes strongly associated with BC^28^. The second OncoOmics approach was made up with the top 40 genes with the highest degree centrality and consensus scoring, such as *TP53*, *ESR1*, *CCND1*, *ERBB2*, *PTEN*, *CDKN1B*, *ATM*, *AKT1*, *STAT3*, *CDH1* and *EGFR*.

The third OncoOmics approach was related to the BC proteome. More than 500 proteins have been identified as strongly involved in oncogenesis. Loss of expression, overexpression or expression of dysfunctional proteins contribute to uncontrolled tumor growth, causing chromosomal rearrangements, gene amplification and ungoverned methylation^59^. Regarding our 230 proteins, 43 showed significant high and low expression (p < 0.001), according to TCGA. The top ten proteins with the highest expression levels were *ERBB2*, *SERPINE2*, *CDH2*, *CCND1*, *EGFR*, *ERCC1*, *IRS1*, *NOTCH1*, *ERBB3* and *INPP4B*, whereas the top ten proteins with the lowest expression levels were *CDH1*, *ATM*, *JAK2*, *MAPK1*, *AKT1*, *AKT3*, *MAPK14*, *ABL1*, *CTNNB1* and *IRF1*. On the other hand, the HPA has analyzed 202 of 230 proteins, where *FOXA1*, *TOP2A*, *CDK2*, *CYP2D6*, *NCOR1* and *RRM1* were involved in oncogenic processes, and *RAC1*, *GJB2*, *MED1*, *PIK3CA*, *PIK3R3*, *FGFR2*, *HCFC2*, *MAP2K4*, *NQO2* and *RAC3* were involved in tumor suppression processes. Lastly, genes with unfavorable prognosis in BC were *RAD51*, *PERP* and *MORC4* (Figure 6)^55,56^. The compendium of all these 60 proteins with significant high and low expression made up the third OncoOmics approach.

The fourth OncoOmics approach was related to the BC dependency map. According to Tsherniak *et al*., the mutations that trigger the growth of cancer cells also confer specific vulnerabilities that normal cells lack, and these dependencies are compelling therapeutic targets^82^. The cancer dependency map identifies essential genes in proliferation and survival of well-annotated cell lines through systematic loss-of-function screens^18–21^. On the one hand, DETEMER2 analyzed the genome-scale RNAi loss-of-function screens. The top 10 genes with the greatest number of significant dependency scores in BC cell lines were *RPL5*, *SF3B1*, *RPA1*, *RRM1*, *BUB1B*, *RPA3*, *RAD51*, *PPP2R1A*, *CHD4* and *POLE*. On the other hand, CERES analyzed the genome-scale CRISPR-Cas9 loss-of-function screens. The top 10 genes with the greatest number of significant dependencies in BC cell lines were *RPA1*, *RRM1*, *TOP2A*, *BUB1B*, *CTCF*, *POLE*, *SF3B1*, *RPL5*, *CCND1* and *SOD2* (Figure 7A). Additionally, the fourth OncoOmics approach was made up of genes with significant dependencies in BC cell lines and all cancer cell lines. *PLCG1*, *CDK4*, *KRAS*, *SPOP*, *CTNNB1*, *EGFR*, *AKT1*, *JAK2*, *MDM4*, *FGFR1*, *IRS1*, *BCL2*, *RELA*, *GATA3*, *PIK3CA*, *PIK3RE*, *PIK3CB*, *FOXA1*, *ERBB3*, *FGFR2*, *ESR1* and *ERBB2* were strongly selective genes, whereas *CDH4*, *TOP2A*, *GNL3*, *RBBP8*, *TOP3A*, *SMARCB1*, *UROD*, *RPL5*, *RAD51*, *PDPK1*, *CCNK*, *SF3B1*, *CDC42*, *ERCC2*, *BUB1B*, *CTCF*, *MAX*, *CCND1*, *BARD1*, *RAC1*, *RPA3*, *SMARCE1*, *PPP2R1A*, *POLE*, *RPA1* and *GRB2* were common essential genes, and *SOD2*, *CDK2*, *ATR*, *RRM1* and *MED1* were both (Figure 7E).

Subsequently, the compendium of the most relevant genes per OncoOmics approach reveals the 144 OncoOmics essential genes in BC (Figure 8A). *RAC1*, *AKT1*, *CCND1*, *PIK3CA* and *ERBB2* were relevant genes in all OncoOmics approaches; *CDH1*, *MAPK14*, *TP53*, *MAPK1*, *SRC* and *RAC3* were relevant genes in the OncoPrint, networking and protein expression analyses; *PLCG1* and *GJB2* were relevant genes in the OncoPrint, networking and DepMap analyses; *MED1*, *TOP2A* and *GATA3* were relevant genes in the DepMap, OncoPrint and protein expression analyses; and *BCL2*, *CTNNB1*, *EGFR* and *CDK2* were relevant in the DepMap, networking and protein expression analyses. Lastly, the top 10 genes with the greatest number of interactions with the hallmarks of cancer were *TP53*, *CTNNB1*, *PTEN*, *KRAS*, *AKT1*, *RAC1*, *EGFR*, *ABL1*, *RB1* and *NOTCH1* (Figure 8D).

According to Reimand *et al*., g:Profiler lets us know the enrichment map of the 144 OncoOmics essential genes in BC^83^. The most significant GO: biological process was the positive regulation of macromolecule metabolic process, the GO: molecular function was phosphatidylinositol 3-kinase activity, the Reactome pathway was generic transcriptor pathway, and the most significant Human Phenotype Ontology term was breast carcinoma^69^. Subsequently, the most relevant network interactions of the GO: biological process and the Reactome pathways were related to immune system, tyrosine kinase, cell cycle and DNA repair terms (Figures 9 and S2)^52,67^.

There is currently great enthusiasm about immunotherapeutic strategies to treat BC. The first approval of an immune checkpoint blockade agent for treatment of BC came in March 2019 when the anti-PD-L1 antibody atezolizumab was approved to be used in combination with nab-paclitaxel for patients with triple-negative BC^84^. 17 OncoOmics essential genes were associated with immunotherapy^62^. Kinases have been recognized as highly tractable targets for BC treatment due to their druggability and critical roles they play in regulating cellular migration, differentiation, growth and survival^85^. 17 OncoOmics essential genes in BC were kinome genes^63^. The cell cycle comprises a series of tightly controlled events that drive cell division and the DNA replication^86^. 12 OncoOmics essential genes in BC were involved in cell cycle^64^. DNA repair constitutes several signaling pathways working in concert to eliminate DNA lesions and maintain genome stability. Defective components in DNA repair machinery are an underlying cause for the development of BC^87^. 19 OncoOmics essential genes in BC were involved in the DNA repair system^65^. RBPs are key players in post-transcriptional events^88,89^. Three recent reports using high-throughput bioinformatics profiling of thousands of tumors now reveal a consistent pattern of alterations in RBPs expression levels across different cancer types^90–92^. Lastly, 11 OncoOmics essential genes were RBPs (Figure 8C)^66^.

Precision medicine provides BC patients with the most appropriate diagnostics and targeted therapies based on the omics profile and other predictive and prognostic tests. Additionally, it is relevant to know the composition of their breast tissue, tumor microenvironment, comorbid conditions and lifestyle^93^.

The OTP is an available resource for the integration of genetics, omics and chemical data to aid systematic drug target identification and prioritization^70^. Currently, there are 111 drugs that are being analyzed in 3151 clinical trials in 39 of the 230 genes. Most of clinical trials are in phase 2; most of the analyzed drugs are small molecules; and most of target classes belong to tyrosine kinases. Finally, the top ten genes with the greatest number of clinical trials in process or completed are *TUBB1*, *ERBB2*, *ESR1*, *TOP2A*, *EGFR*, *ESR2*, *VEGFA*, *CDK4*, *POLE* and *RRM1*^70^ (Figure 10).

PharmGKB collects the precise guidelines for the application of pharmacogenomics in clinical practice^41–44^. This database details 154 clinical annotations associated with 70 genes in BC. The CGI is a platform that annotates clinical evidence and tumor variants that constitute state-of-art biomarkers of drug response. The CGI details 76 clinical annotations associated with 26 genes in BC^71^. According to TCGA, PCA details 648 clinical annotations associated with 14 genes in BC^72^. Lastly, the drug-gene interaction matrix is a compendium of the most relevant clinical annotations made up of 32 genes and 51 drugs in order to facilitate the treatment of patients with BC (Figure S3).

In conclusion, since BC is a complex and heterogeneous disease, the study of different OncoOmics approaches is an effective way to reveal essential genes to better understand the molecular landscape of processes behind oncogenesis, and to develop better therapeutic treatments focused on pharmacogenomics and precision medicine.

## METHODS

### OncoPrint of genetic alterations according to the Pan-Cancer Atlas

PCA has reported the clinical data of 1084 individuals with BC and it can be visualized in the Genomic Data Commons of the National Cancer Institute (https://gdc.cancer.gov/) and in the cBioPortal (http://www.cbioportal.org/)^45,46^. The clinical annotations were age, pTNM classification, tumor type, tumor stage and race/ethnicity.

Additionally, PCA has reported genetic alterations (mRNA upregulation, mRNA downregulation, CNV amplification, CVN deep deletion, missense mutation, truncating mutation, inframe mutation and fusion gene) in 994 individuals. Putative mutations were analyzed through exome sequencing, CNVs through the Genomic Identification of Significant Targets in Cancer (GISTIC 2.0)^94,95^, and mRNA expression through RNA Seq V2. We analyzed five gene sets in order to compare the average frequency of genetic alterations among them. The first gene set (n = 177) was integrated by the non-cancer genes^96^. We calculated the OncoScore of non-cancer genes, taking out all genes from our study. The second gene set (n = 119) was the BC driver genes, according to The Network of Cancer Genes^61^. The third gene set (n = 84) was taken from our previous study where we developed a Consensus Strategy of prioritized genes related to BC pathogenesis^28^. The fourth gene set (n = 85) was made up of genes associated with BC development, according to several PCA studies^30,31,57^. The fifth gene set (n = 91) consisted of BC biomarkers and druggable enzymes taken from PharmGKB and the CGI (Table S2)^37,38,40^. Finally, the significant differentiation of the average frequency of genetic alterations among gene sets was analyzed (p-value < 0.001).

The OncoOmics approaches were performed in 230 genes conformed by the CS, PCA and PharmGKB/CGI gene sets. Firstly, we calculated the percentage and ratio of genetic alterations per intrinsic molecular subtype and tumor stage, and we established a ranking of genes with the greatest number of different genetic alterations. Subsequently, we performed an OncoPrint of genes with more genetic alterations than the average. The final list of genes made up the first OncoOmics approach.

### Pathway enrichment analysis

The enrichment analysis of signaling pathways was performed using David Bioinformatics Resource to obtain integrated information from KEGG^47–50^. It was carried on in the 230 genes, taking into account terms with a significant FDR < 0.01. After that, genetic alterations that comprise each signaling pathway were analyzed, taking into account the molecular subtype and tumor stage of individuals from PCA. Circos plots and violin plots were designed to visualize all data. Lastly, in order to compare the ratio of genetic alterations among subtypes and tumor stages, normalization was carried out dividing the number of genetic alterations by the number of individuals per subtype and tumor stage. Regarding molecular subtypes, 499 individuals were luminal A, 197 were luminal B, 171 were basal-like, 78 were Her2-enriched and 36 were normal-like, and regarding tumor stage, 255 were stage T1, 586 were stage T2, 113 were stage T3 and 103 were stage T4.

### Protein-protein interaction network

The PPi network with a highest confidence cutoff of 0.9 and zero node addition was created using the String Database, which takes into account predicted and known interactions^51^. The confidence scoring is the approximate probability that a predicted link exists between two enzymes in the same metabolic map, whereas the degree centrality of a node means the number of edges the node has to other nodes in a network. The centrality indexes calculation and network visualization were analyzed through the Cytoscape software^52^. Genes with the highest degree centrality, consensus score and sub-networks were differentiated by colors in the PPi network. On the other hand, OncoPPi (http://oncoppi.emory.edu/) reports the development of a cancer-focused PPi network, identifying more than 260 high-confidence cancer-associated PPi^53,54^. In addition, the OncoPPi BC network consisted of 16 genes and 18 PPi experimentally analyzed in BC cell lines^53,54^. The correlation of the degree centrality by means of Spearman p-value test between our String PPi network and the OncoPPi BC network allowed for the validation of all the high-confidence BC-focused PPi analyzed in cell lines^28^. Lastly, genes with the highest degree centrality and consensus scoring made up the second OncoOmics approach.

### Protein expression analysis

TCGA has reported the protein expression data of 994 individuals with BC through RPPA and mass spectrometry by the Clinical Proteomic Tumor Analysis Consortium (CPTAC), and it can be visualized in the cBioPortal^45,46^. We analyzed the protein expression of 230 genes (CS, PCA and PharmGKB/CGI gene sets) where Z-scores _≥_ 2 mean a significant high protein expression and Z-scores _≤_ −2 mean a significant low protein expression. On the other hand, the Human Protein Atlas (https://www.proteinatlas.org/) explains the diverse molecular signatures of proteomes in the human tissues based on an integrated omics approach that involves quantitative transcriptomics and tissue microarray-based immunohistochemistry^56,58,59^. We compared the protein gene levels (high, medium, low and non-detected) of our 230 genes between normal and BC tissues. Finally, we analyzed the overall survival curve of our 230 genes and reveled all biomarkers with significant unfavorable prognostic (p < 0.001)^55,56^. All genes with the altered protein expression made up the third OncoOmics approach.

### Breast cancer dependency map

The DepMap project (https://depmap.org/portal/) is a collaboration between the Broad Institute and the Welcome Sanger Institute. Multiple genetic or epigenetic changes provide cancer cells with specific vulnerabilities that normal cells lack. Even though the landscape of genetic alterations has been extensively studied to date, we have limited understanding of the biological impact of these alterations in the development of specific tumor vulnerabilities, which triggers a limited use of precision medicine in the clinical practice worldwide. Therefore, the main goal of DepMap is to create a comprehensive preclinical reference map connecting tumor features with tumor dependencies to accelerate the development of precision treatments^18–21^.

In order to identify essential genes for BC cell proliferation and survival, DepMap performed systematic loss-of-function screens in a large number of well-annotated BC cell lines representing the tumor heterogeneity and their molecular subtypes. The DEMETER2 algorithm was applied to analyze genome-scale RNAi loss-of-function screens in 73 BC cell lines and 711 cancer cell lines, whereas the CERES algorithm was applied to analyze genome-scale CRISPR-Cas9 loss-of-function screens in 28 BC cell lines and 558 cancer cell lines^19,21^. In addition to existing cell lines, the Cancer Cell Line Encyclopedia (CCLE) project will greatly expand the collection of characterized cell lines to improve precision treatments^97^.

Regarding dependency scores, a lower score means that a gene is more likely to be dependent in a specific cancer cell line. A score of 0 means that a gene is not essential, whereas a score of −1 corresponds to the median of all common essential genes. A strongly selective gene means that its dependency is at least 100 times more likely to have been sampled from a skewed distribution than a normal distribution. Lastly, a common essential gene is when in a pan-cancer screen its gene ranks in the top most depleting genes in at least 90% of cell lines^18^. All genes with a dependency score ≤ −1 made up the fourth OncoOmics approach.

### Enrichment map of the OncoOmics essential genes in BC

The pathway enrichment analysis gives scientists curated interpretation of gene lists generated from genome-scale experiments^67^. The OncoOmics essential genes in BC were analyzed by using g:Profiler (https://biit.cs.ut.ee/gprofiler/) in order to obtain significant annotations (FDR < 0.001) related to GO terms, pathways, networks and disease phenotypes. Subsequently, g:Profiler annotations were analyzed with the EnrichmentMap software in order to generate network interactions of the most relevant GO: biological processes and Reactome pathways, and these networks were visualized using Cytoscape^52,67^.

### Precision medicine

We analyzed drug-gene interactions for BC using four selective databases: 1) OTP^70^, 2) PharmGKB^37,40^, 3) CGI^38^, and 4) PCA^98^. The Open Targets Platform (https://www.targetvalidation.org) is comprehensive and robust data integration for access to and visualization of potential drug targets associated with BC. Additionally, this platform shows all drugs in clinical trials associated with BC genes, detailing its phase, status, type and target class^70^. PharmGKB (https://www.pharmgkb.org/) collects complete guidelines for application of pharmacogenomics in clinical practice, according to several consortiums worldwide^41–44^. The CGI (https://www.cancergenomeinterpreter.org/home) flags genomic biomarkers of drug response with different levels of clinical relevance^38^. Finally, PCA reveals genetic alterations, druggable enzymes and clinical annotations in a cohort of 994 individuals^3,12,30–36^. The clinical annotations of these four databases were analyzed in order to create a drug-gene interaction matrix.

## Acknowledgments

This work was supported by Universidad UTE (Quito, Ecuador), Universidad de Las Américas (Quito, Ecuador), University of Coruna (Coruña, Spain), University of the Basque Country (Bilbao, Spain), and McGill University (Montreal, Canada). Additionally, this work was supported by “Collaborative Project in Genomic Data Integration (CICLOGEN)” PI17/01826 funded by the Carlos III Health Institute from the Spanish National plan for Scientific and Technical Research and Innovation 2013-2016 and the European Regional Development Funds (FEDER).

## Author Contributions

ALC and ET conceived the subject and the conceptualization of the study. ALC wrote the manuscript. ET, SJB, CRM, HGD and CPyM supervised the project. ALC and CPyM did founding acquisition. ALC, SG and ACA did data curation and supplementary data. ET, SG, ACA, SJB, CRM, HGD, AP, YPC and CPyM gave conceptual advice and valuable scientific input. Finally, all authors reviewed the manuscript.

## Data availability statement

All data generated or analysed during this study are included in this published article (and its Supplementary Information files).

