## Supplementary figures and images for "OncoOmics approaches to reveal essential genes in breast cancer: a panoramic view from pathogenesis to precision medicine"

### Supplementary Figure 1

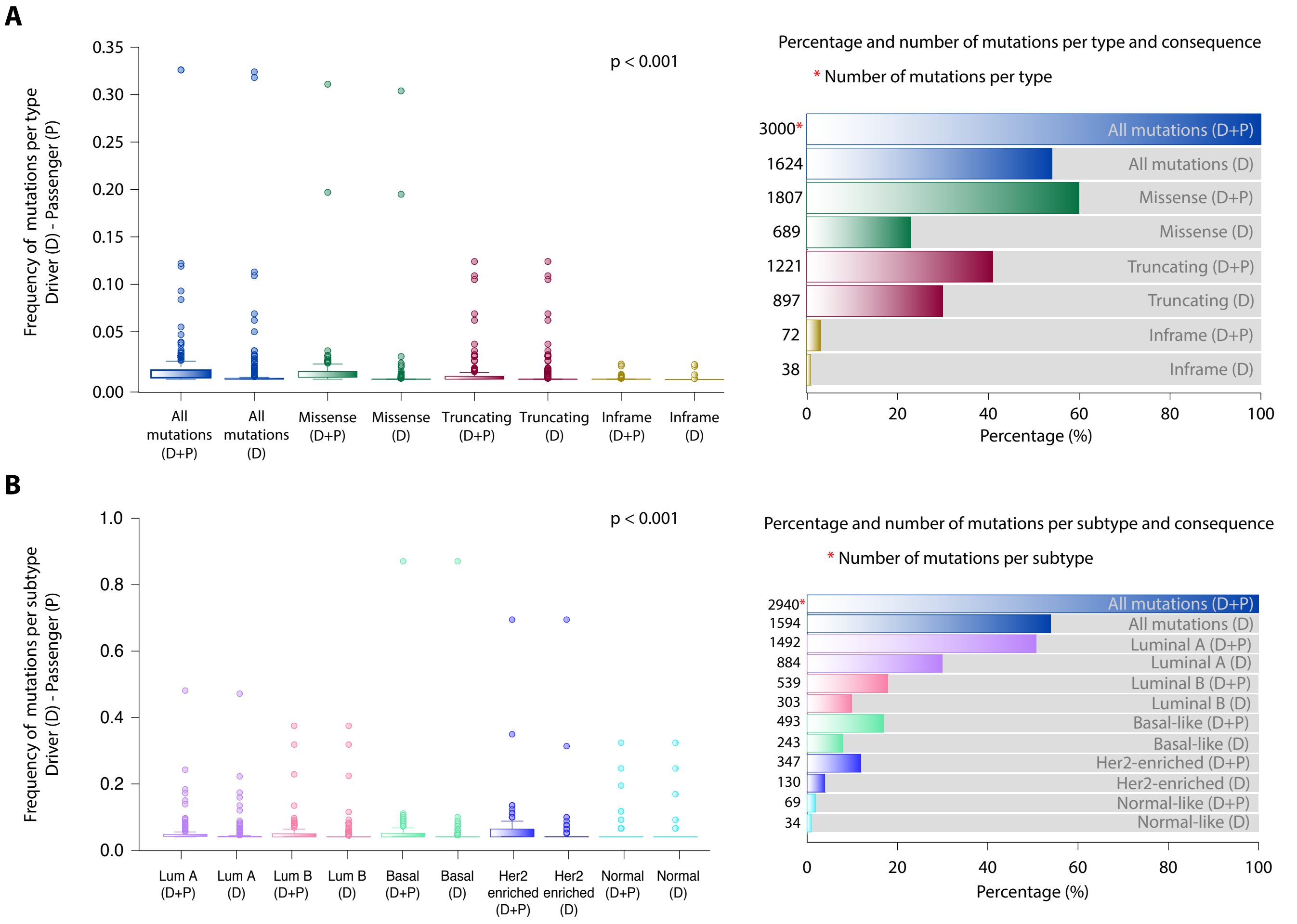

### Supplementary Figure 2

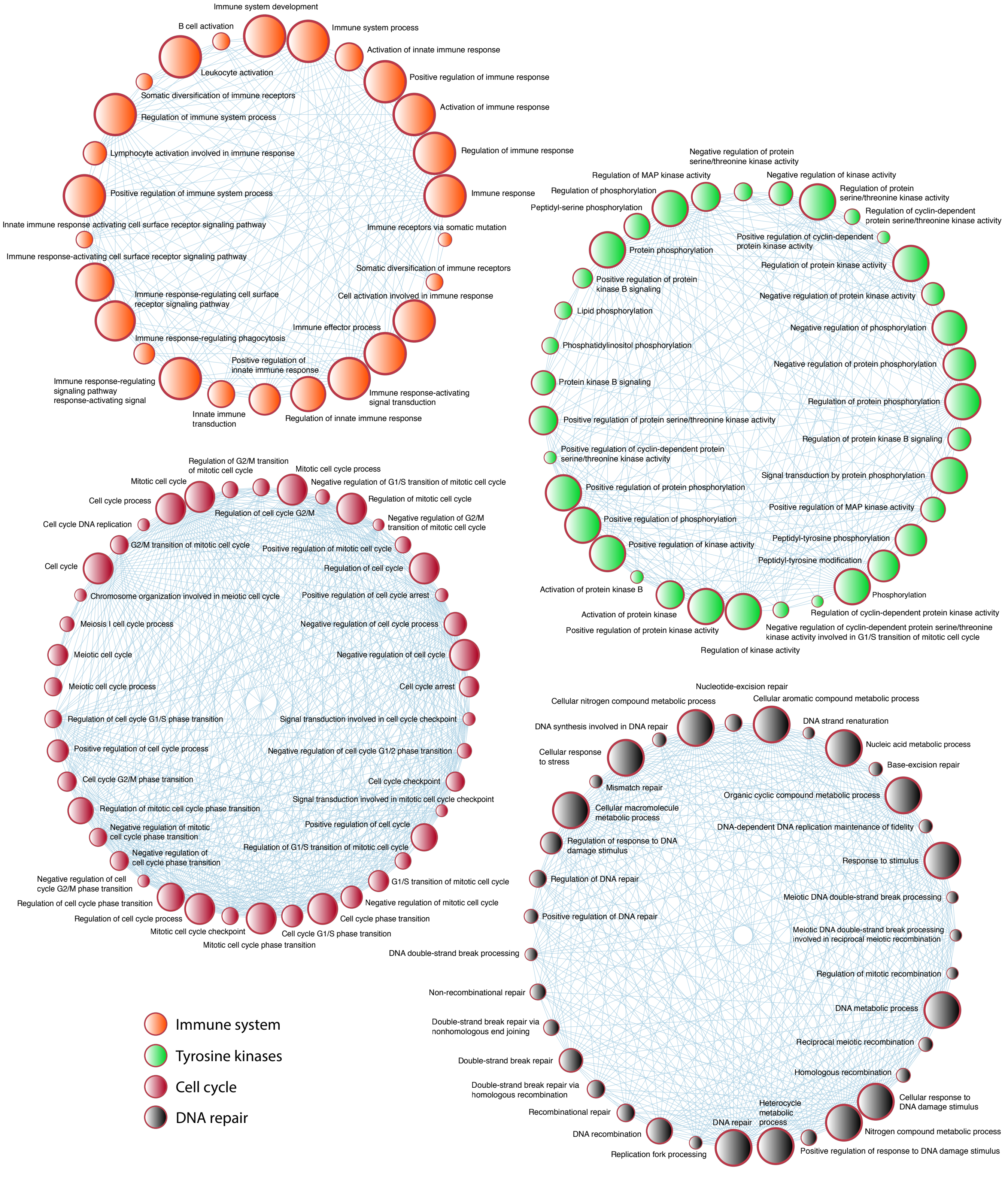

### Supplementary Figure 3

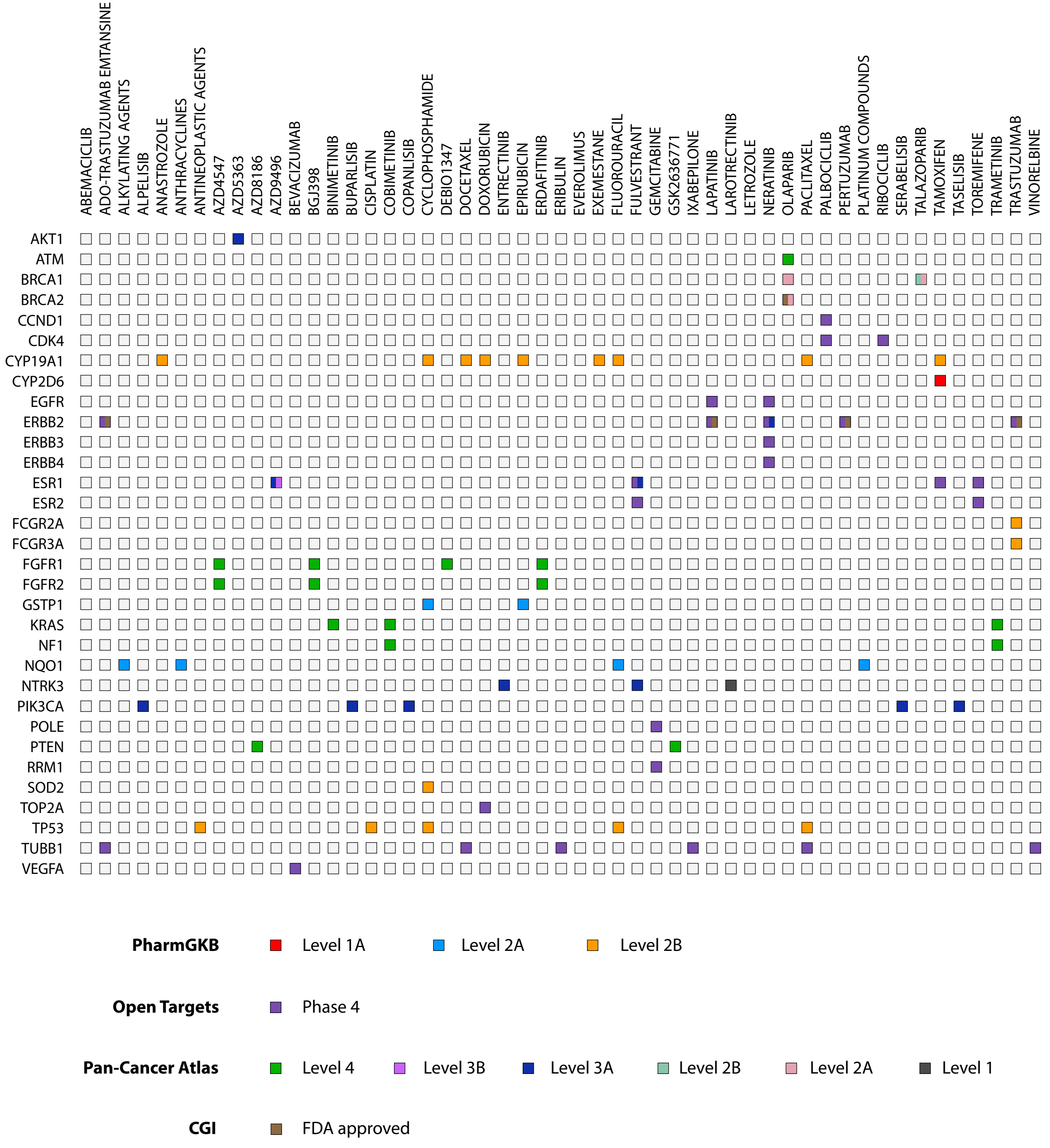
